# Effects of alcohol consumption and tobacco smoking on the composition of the ensemble of drug metabolizing enzymes and transporters in human liver

**DOI:** 10.1101/2024.05.14.594255

**Authors:** Kari A. Gaither, Dilip Kumar Singh, Guihua Yue, Julia Trudeau, Kannapiran Ponraj, Nadezhda Y. Davydova, Philip Lazarus, Dmitri R. Davydov, Bhagwat Prasad

## Abstract

We examined the effect of alcohol consumption and smoking on the abundance of drug-metabolizing enzymes and transporters (DMET) in human liver microsomes (HLM) isolated from liver tissues of 94 donors. Global proteomics analysis was performed and DMET protein levels were analyzed in relation to alcohol consumption levels, smoking history, and sex using non-parametric tests (p-value ≤ 0.05; cutoff of 1.25-fold change, FC). The examination of the alcohol-induced changes was further enforced by correlational analysis, where we used arbitrary alcohol consumption grade (ACG) scaling from 0 to 4 to establish a set of protein markers. We elaborated a provisional index of alcohol exposure (PIAE) based on a combination of relative abundances of four proteins (ER chaperone HSPA5, protein disulfide isomerases PDIA3 and P4HB, and cocaine esterase CES2) best correlating with ACG. The PIAE index was then used to find its correlations with the abundances of DMET proteins. Our results demonstrate considerable alcohol-induced changes in composition of the pool of cytochrome P450 enzymes in HLM. We observed significantly increased abundances of CYP2E1, CYP2B6, CYP2J2, and NADPH-cytochrome P450 reductase. In contrast, CYP1A2, CYP2C8, CYP2C9, CYP4A11, and cytochrome b_5_ protein levels were downregulated. Significant alteration in abundances of UDP-glucuronosyltransferase (UGT) were also detected, comprising of elevated UGT1A6, UGT1A9, and UGT2A1, and reduced UGT1A3, UGT1A4, UGT2B7, UGT2B10, and UGT2B15 levels. Important alcohol-induced changes were also observed in the expression of non-CYP and non-UGT DMET. Additionally, tobacco smoke was associated with elevated CYP1A2, UGT1A6, UGT2A1, and UGT2B4 and decreased FMO3, FMO4, and FMO5 levels.

## 1. Introduction

Alcohol and tobacco are widely used substances worldwide and are associated with numerous health risks. In 2021, 60 million people in the US reported drinking alcohol excessively and over 11 million smoked at least a pack of cigarettes per day (SAMHSA, 2022). Together, they ranked in the top ten risk factors for global disease burden and resulted in over 10 million combined deaths in 2019 (Shield et al., 2020; G. B. D. Tobacco Collaborators, 2021). An important part of the alcohol-related fatalities is associated with deleterious alcohol interactions with drugs, including both prescribed medications and the drugs of abuse. There are numerous known examples of considerable changes in drug pharmacokinetics and pharmacodynamics by both chronic and acute alcohol exposure (Djordjevic et al., 1998; Weathermon and Crabb, 1999; Meskar et al., 2001; Tanaka, 2003; Tom, 2008; Chan and Anderson, 2014). Consideration and prediction of alcohol-drug interactions (ADI) are crucial for practical pharmacotherapy, as they may dramatically affect the optimal dosage of drugs in alcohol consumers. Life-threatening ADI with acute alcohol consumption, such as those observed with benzodiazepines, tricyclic antidepressants, opiates, and barbiturates, are revealed in the cases of fatal co-intoxications (Koski et al., 2003; Koski et al., 2005; Gomes et al., 2017). In contrast to these easy-to-recognize acute occurrences, many instances of drug interactions with chronic alcohol exposure may remain unidentified.

However, our understanding of the mechanisms of the effects of alcohol on drug metabolism remains limited. Although the significant increase in cytochrome P450 2E1 (CYP2E1) content in the liver and other tissues observed in both alcoholics and moderate alcohol consumers represents one of the most important effects of alcohol on protein expression (Cederbaum, 1998; Cederbaum, 2006), its involvement in the instances of ADI is commonly considered insignificant due to its minor role in drug metabolism (Jang and Harris, 2007; Chan and Anderson, 2014) with the well-known exception of acetaminophen (Lieber, 1999). However, the impacts of the induction of CYP2E1 by alcohol on drug metabolism and other functions of the cytochrome P450 enzymes (P450s) appear to be underestimated. For example, interactions of CYP2E1 with other P450s provides the most likely explanation for the alcohol-induced increase in the metabolism of CYP3A substrates diazepam and doxycycline (Sellman et al., 1975; Neuvonen et al., 1976), or phenytoin, tolbu tamide, and warfarin (Kater et al., 1969a; Sandor et al., 1981) metabolized primarily by CYP2C9. Evidence of a direct cause-to-effect relationship between alcohol-dependent induction of CYP2E1 and the effects of this kind is provided by our studies of the impact of CYP2E1 on the activity of CYP3A4, CYP1A2, and CYP2C19 (Davydova et al., 2019; Dangi et al., 2021).

Of note, tobacco smoke is comprised of over 7000 chemicals at least 69 of which are known carcinogens (Hoffmann et al., 2001; Soleimani et al., 2022). Upon entering the lung, chemicals in tobacco smoke make their way to the liver where they are metabolized, interact with, and in many cases induce the expression of drug metabolizing enzymes (O’Malley et al., 2014). Thus, various drug interactions may occur with tobacco users (Maideen, 2019). Sex is also known to contribute to differences in drug m etabolizing enzymes, for instance recent findings indicate a 2.6 fold greater level of UDP-glucuronosyltransferase (UGT) 2B17, critically involved in the metabolism of several pharmaceuticals, in males than females (Bhatt et al., 2018), but this is an understudied area overall.

The goal of the present study was to investigate the impact of alcohol consumption and smoking tobacco on the abundance of drug-metabolizing and transporter (DMET) proteins in human liver, using a large series of human liver microsomes (HLM) from donors with reported history of alcohol consumption and tobacco smoking. A secondary goal was to investigate any differences in the expression of drug metabolizing enzymes and transporter proteins due to sex. Using high throughput quantitative proteomics method, we quantified DMET proteins present in HLM isolated from liver tissues from 94 adult donors (61 males and 33 females) with documented alcohol intake and smoking histories. Large-scale studies of the DMET proteome in postmortem liver samples are few, typically low in sample size, and particularly lacking information about the effects of sex and lifestyle factors with detailed histories (Wegler et al., 2022; Pedersen et al., 2023). Our findings will contribute to a clinically translational understanding of changes in the levels of proteins involved in drug metabolism and distribution with a look to the future of enabling optimal dosing for precision medicine.

## 2. Materials and Methods

### 2.1. Chemicals and reagents

Liquid chromatography−mass spectrometry (LC−MS)-grade acetonitrile, methanol, chloroform, and formic acid were procured from Fisher Scientific (Fair Lawn, NJ) while acetone was obtained from Sigma-Aldrich (St. Louis, MO). Ammonium bicarbonate (98% pure), dithiothreitol, iodoacetamide, and MS-grade trypsin were purchased from Thermo Fisher Scientific (Rockford, IL). The bicinchoninic acid (BCA) kit was from Pierce Biotechnology (Rockford, IL).

### 2.2 Human liver microsomes

The majority of the liver samples used in this study (n=88) were from the biobanks established in the Prasad and Lazarus Laboratories. Additionally, six liver tissue samples from moderate-to-heavy alcohol consumers were procured from BioIVT corporation (Westbury, NY ). The base inclusion criterion was a documented alcohol intake history. One aim of selection was to identify and include donors with a history of heavy alcohol consumption while excluding the use of illicit substances as potential confounders where possible. Microsomes were prepared from 71 liver specimens from the Lazarus Laboratory via differential centrifugation as described previously (Jones et al., 2012). Preparation of microsomal fractions from 17 liver specimens was performed as described by Nelson et al. with minor modifications (Nelson et al., 2001). Frozen liver tissue specimens (0.5 - 1 g) were covered with 0.1 M potassium phosphate buffer, pH 7.4, containing 0.125 M KCl, 0.25 M sucrose, and 1.0 mM EDTA (Buffer A) at room temperature and allowed to thaw. After decanting the buffer, the tissue was minced with scissors on ice and supplemented with 2.5 volumes of ice-cold buffer A containing 0.25 mM PMSF. The mixture was homogenized on ice with 10 strokes in a glass homogenizer with a motorized Teflon pestle. After diluting the homogenate to 7 - 8 volumes of the sample weight with ice-cold PMSF-containing buffer A, it was centrifuged at 10,500 *x*g for 40 min. The pellet was discarded, and the supernatant was centrifuged at 118,000 *x*g for 90 min. The upper lipid layer and the supernatant were discarded. The pellet was resuspended in 0.1 M Na-HEPES buffer containing 60mM KCl and 0.25 M sucrose, pH 7.4 (Buffer B) added to reach 1.5 - 2 volumes of the sample weight and centrifuged at 118,000 *x*g for 90 min. The pellet was resuspended in Buffer B (1 ml per 1 g of tissue) with a syringe and plastic pestle in a 1.5 ml Eppendorf tube and stored at -80 °C.

### 2.3. Trypsin digestion and sample preparation for proteomics analysis

HLM were analyzed for protein content using a BCA kit via standard vendor protocols and then digested by an optimized trypsin digestion protocol described previously (Sharma et al., 2023). Briefly, 1 mg/ml protein samples (80 μg) in 100 mM ammonium bicarbonate, pH 7.8, were reduced and denatured by the addition of 250 mM dithiothreitol and incubation at 95°C for 10 min under gentle shaking. After samples were brought to room temperature, proteins were then alkylated via the addition of 100 mM iodoacetamide (incubated in the dark for 30 min). Protein precipitation was brought about by the addition of ice-cold acetone at -80 °C for 1 hour. Samples were then centrifuged at 16,000 *x*g for 10 minutes at 4 °C and the supernatant was discarded. Samples were washed with ice-cold methanol followed by another round of high-speed centrifugation at 16,000 *x*g for 10 minutes at 4 °C. The supernatant was removed, and the pellet was dried for 30 min at room temperature. The pellet was then resuspended in 60 μl ammonium bicarbonate buffer (50 mM, pH 7.8). Protein digestion occurred via the addition of 20 μl trypsin (50:1, protein: trypsin ratio) with gentle shaking at 37 °C for 16 hours and was quenched with 5 μL of 5% formic acid in water. The sample was then centrifuged at 16,000 *x*g for 10 minutes at 4 °C and stored at −80 °C until LC-MS analysis.

### 2.3. LC-MS data acquisition

Global quantitative proteomics analysis was performed using the Easy Spray 1200 series nanoLC coupled with Orbitrap Q-Exactive Mass Spectrometer (Thermo Fisher Scientific, Waltham, MA). 1 μl of protein digest sample (1 μg/μl) was injected and peptides separation was achieved using a Thermo Scientific Acclaim Pepmap RSLC C18 25 cm x 75 μm (2 μm, 100 Å) column with a mobile phase consisted of 0.1% formic acid in water (A) and 80% acetonitrile with 0.1 % formic acid (B). The flow rate was 300 nl/min with a 35 min gradient as follows: 0-2 min (0-10% B), 2-27 min (10-45% B), 27-28 min (45-100% B), 28-35 min (100% B).

The eluted peptides were detected in DIA (data-independent acquisition) mode. The spray voltage was 1.7 kV with 300 °C capillary heat. The MS1 scan range was set to m/z 348–1100 with a mass resolution of 60,000, an auto gain control (AGC) target of 3 × 10^6^, and a maximum injection time of 55 msec. MS2 was set to a resolution of 30,000, AGC of 1 × 10^6^, normalized collision energy of 30, and a maximum injection time of 55 msec. The DIA had a variable isolation window set to 25 m/z spanning the 350-400 mass range, 20 m/z spanning the 400-870 mass range, and 40 m/z spanning the 870-1110 mass range.

### 2.4. Proteomics data analysis

DIA-NN (version 18.1.1) (https://github.com/vdemichev/DiaNN), was used for library-free analysis (Demichev et al., 2020). Deep learning-based in silico spectral library generation was enabled with the human FASTA database. The identified peptides had a maximum number of missed cleavages set to 1 as the default. Fixed modification includes carbamidomethylation, N-terminal methionine excision, and variable modification including the oxidation of methionine residues and acetylation of protein N termini. Peptides were identified with a 1% false discovery rate. An unrelated run was selected, and all other parameters used the default setting.

To account for batch-to-batch or interlaboratory technical variability during the preparation of subcellular fractions, we normalized DMET proteins to the sum of a set of 75 proteins with known localization in the microsomal membrane or lumen (see Table 1 in Supplementary Materials for the list). For this normalization, the raw MS intensities (MSI) for each (*i*-th) protein in each (*j* -th) HLM sample were first normalized on the total of intensities of all 75 proteins in this sample:

**Table 1.**
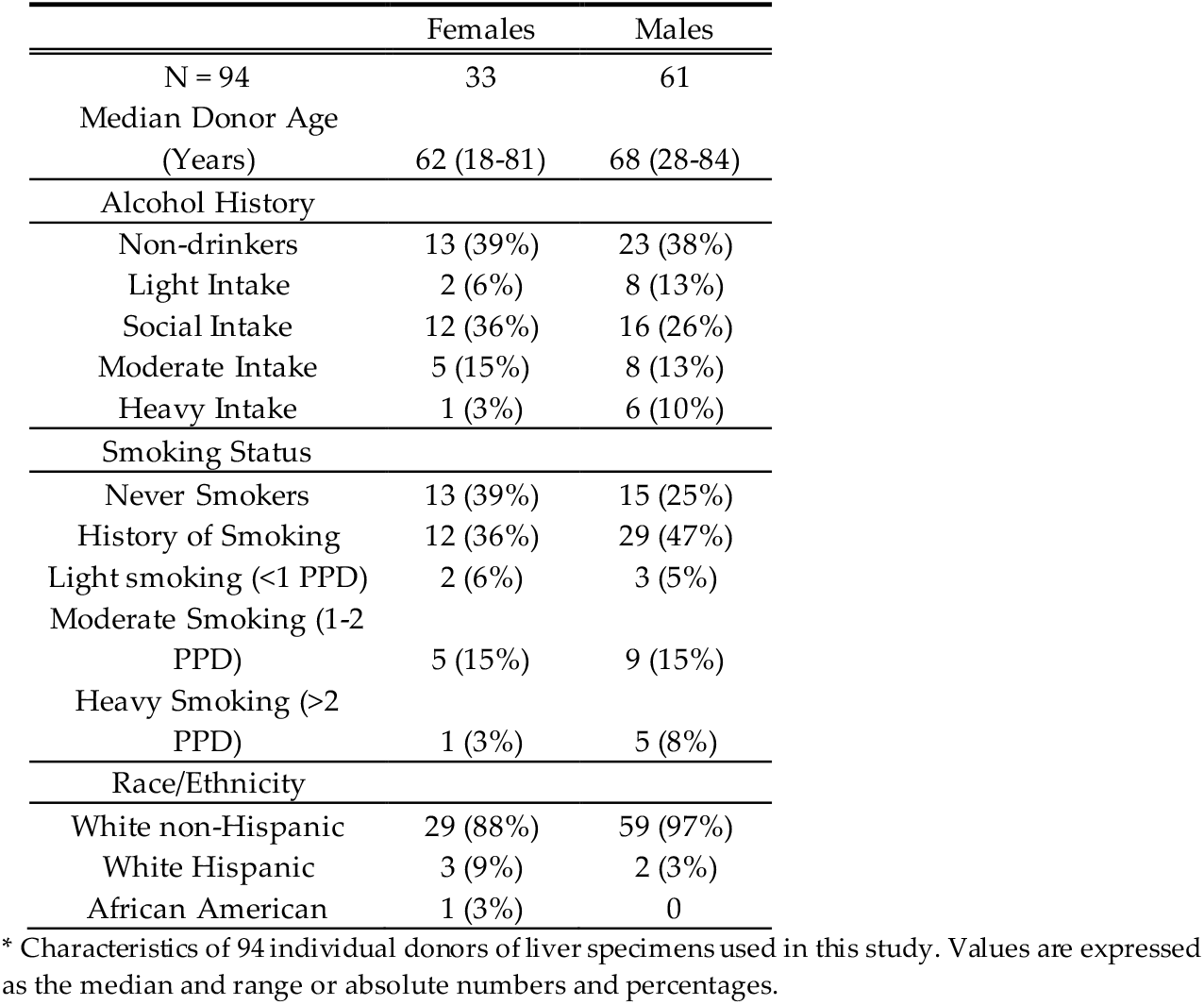
Characteristics of study population*.

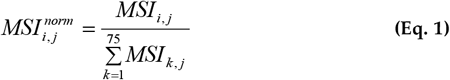

We used the total protein approach (TPA) to compare the abundances of proteins between different treatment groups in a quantitative manner (Wisniewski and Rakus, 2014). This method yields accuracy similar to that obtained by targeted proteomics for protein quantification while allowing for a broader coverage of proteins detected (Sharma et al., 2023). Using this approach makes it possible to determine the protein concentration (mg individual protein per mg total protein), of individual proteins within a sample as follows:

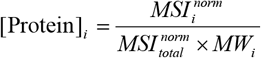

Where 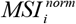 is the normalized MS intensity defined as the normalized (see Eq. 1) sum of the MS1 spectral intensities of all peptides identified to match the sequence of i-th protein, 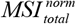 is the sum of all normalized MS spectral intensities for all proteins in a particular sample, and MW_i_ is the molecular weight of the i-th protein. Pathways were determined by using the STRING database (www.string-db.org) using the homo sapiens background (Szklarczyk et al., 2019).

### 2.5. Non-parametric statistical analysis and data visualization

To probe for differences in DMET proteins with alcohol consumption, individual-derived HLM were segregated by alcohol intake history into five groups: (1) non-drinking control group, including those with a past history of drinking, (2) light alcohol drinkers, (3 ) social drinkers, (4) moderate alcohol drinkers, and (5) heavy alcohol drinkers. Light alcohol consumption was defined as 1 alcoholic drink or less per day, moderate as 2-3 drinks per day, and heavy as >3 drinks per day. Social alcohol as a category lacks a definitive quantity consumed and was ordered between light and moderate groups. GraphPad Prism 8.4.3 (GraphPad Software, La Jolla, CA) was used to perform statistical analysis and produce Volcano plots and bar charts. DMET proteins were analyzed for significant differences using the student t-test with Welch’s correction for unequal variance. A p-value of <0.05 was considered significant. BioRender (Toronto, Ontario) was used to produce diagrams and InteractiVenn (Heberle et al., 2015) was used to create VENN diagrams. Pie charts were produced in Microsoft Excel for Microsoft 365 MSO (Version 2402 Build 16.0.17328.20282) 64-bit (Microsoft, Redmond, WA). Correlation analysis was performed using SpectraLab data analysis software (http://cyp3a4.chem.wsu.edu/spectralab.html).

### 2.6. Analysis of correlations of protein abundances with the level of alcohol consumption

In our further analysis of the effects of alcohol exposure on the HLM proteome we focused on a subset of 75 DME- and ER-stress-related proteins with known localization in the microsomal membrane or microsomal lumen (see Table 1 in Supplementary Material).

To assess the level of alcohol exposure of liver donors numerically, we established the Alcohol Consumption Grade (ACG) scale, where the donors indicated as “non-drinkers”, “former drinkers”, “light drinkers”, “social drinkers”, “low-to-moderate drinkers”, “moderate drinkers” and “heavy drinkers” were assigned the grades of 0, 0.5, 1, 1.5, 2, 3 and 4, respectively. These arbitrary assignments were done based on demographic records available for the donors. This gradation slightly differs from the one used in the preliminary analysis. Specifically, we grouped the former drinkers in a separate category as their claim of abstinence may be an exaggeration. We also divided moderate drinkers into two groups, where “low-to-moderate drinkers” were defined as the donors consuming two drinks per day, while “moderate drinkers” were those with three drinks per day consumption.

To find the correlation of the ACG scores with variations of the protein abundances, we calculated the vectors of relative protein abundance (VRA) for each of the 75 proteins. In these calculations, we used the MS intensities normalized on the total of intensities of all 75 selected proteins in this sample (see Eq. 1 above). After normalization, the averaged normalized intensity for each protein in all 94 HLM samples was assessed and used to calculate the VRA value for each (*i*-th) protein in each (*j*-th) HLM samples:

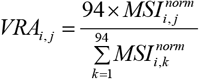

The resulting 75 VRA vectors, which reflect the relative differences in protein abundances between the HLM samples, were used to find their linear combination approximating the vector of Alcohol Consumption Grade. In this search, we approximated the ACG vector by a linear combination of several VRA using the multidimensional linear least-square regression algorithm as described in (Arthurs, 1963). This algorithm was sequentially applied to every possible combination of 2 – 4 proteins to find a numerical scale best reflecting apparent alcohol exposure of the liver donors (Provisional Index of Alcohol Exposure, PIAE). The search algorithm described above was implemented using our SpectraLab data analysis software. The found PIAE vector was then used to probe its correlations with VRA for all 75 selected proteins.

## 3. Results

### 3.1. Liver specimens

HLM from 94 individuals (33 females and 61 males) with known alcohol intake histories were analyzed by LC-MS, with the process outlined in Fig 1A. Specimens were stratified by alcohol intake into the categories of non-drinkers, light drinkers, social alcohol users, moderate drinkers, and heavy drinkers. The set included 69 non-smokers and 20 smokers of >1 pack per day (ppd) (5 were considered light smokers) (Fig 1B). The characteristics of the study population are shown in Table 1. Due to finite resources, heavy alcohol users were limited to 7 individuals, while non-drinking individuals numbered 36. Overall, females were outnumbered almost 1:2 by males with the median donor age similar in females (62 yr) and males (68 yr). Although alcohol intake was the major focus of our analysis, we broadened our efforts to include the impact of smoking and sex differences on the DMET proteome.

**Fig. 1.**
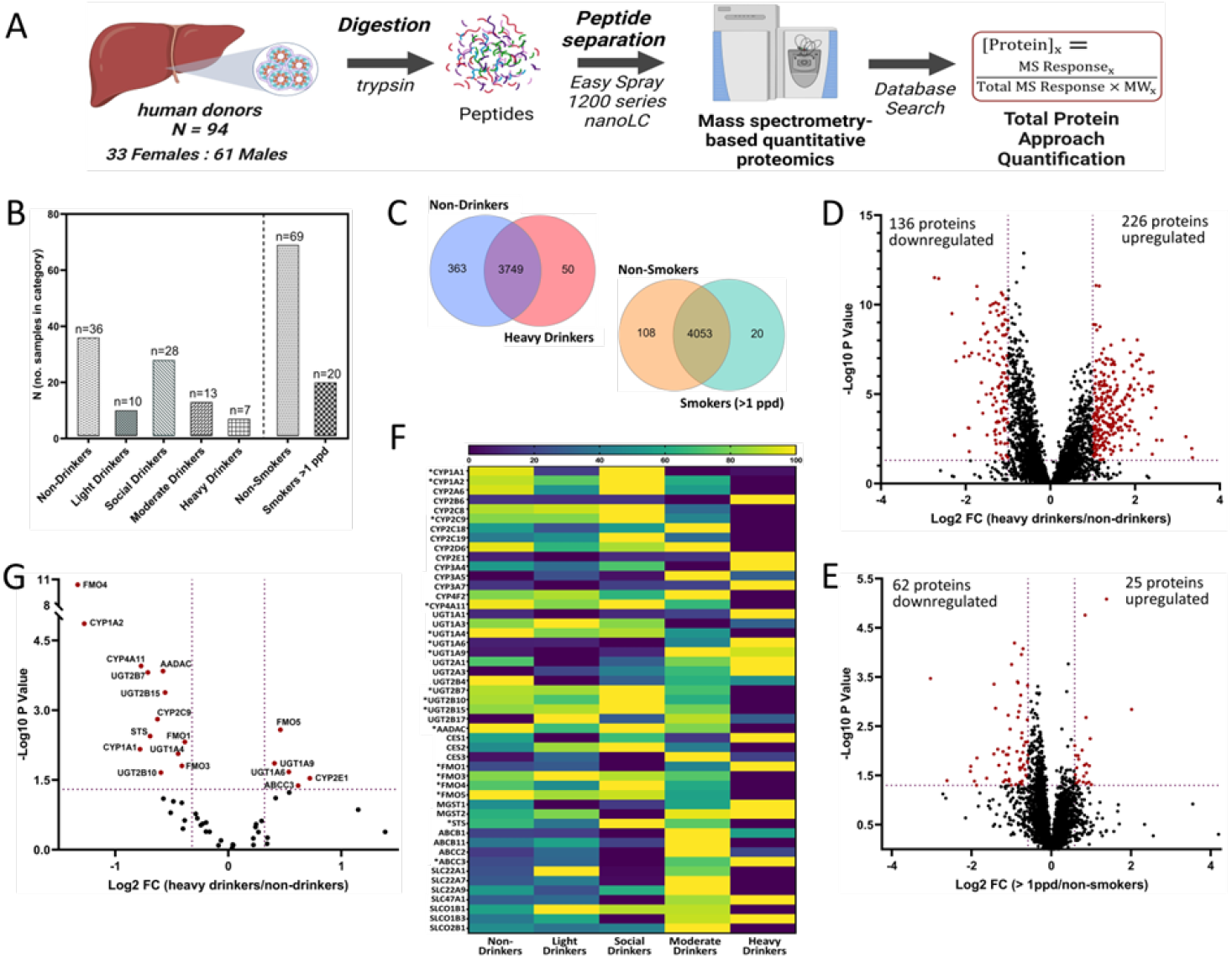
Global proteomics analysis of human liver microsomes prepared from 94 individuals with known alcohol intake histories. A) Mass spectrometry-based workflow scheme of human liver microsomes (N=94, 33 females and 61 males). B) HLM specimens from individuals in each alcohol and smoking history category. C) Venn Diagram of proteins detected and overlapping in non-drinkers vs heavy drinkers and individuals who smoked > 1ppd vs non-smokers. D) Differentially expressed proteins in HLM derived from individuals with heavy alcohol intake compared to non-drinking individuals (P<0.05, Fold change cut off of 2.0. E) Differentially expressed proteins in HLM derived from individuals who smoked > 1ppd compared to non-smoking individuals (P<0.05, Fold change cut off of 1.5. F) Heat map of all DMET proteins resulting from label-free quantitative proteomic analysis of individual-derived HLM, based on the normalized mean protein levels for each DMET across the alcohol use categories. Proteins with significantly different levels of expression in the HLM from heavy drinkers as compared to non-drinkers are indicated with an asterisk (P value < 0.05). G) Differential expression of DMET proteins in individual-derived HLM from heavy drinkers as compared to non-drinkers. (P value <0.05, fold-change cutoff of 1.25).

### 3.1. Effects of alcohol consumption and tobacco smoke on global proteome

In total, 4198 proteins were detected across all groups. A comparison of the global proteomics data for the non-drinking control group and the heavy drinking group yielded 4162 proteins with the majority overlapping; over 360 proteins were unique to HLM from non-drinkers and 50 were unique to HLM from heavy drinkers (Fig 1C). Conversely when the non-smoking group and the >1ppd smoking group were compared, only 128 proteins out of 4181 detected were unique across the 2 groups; 108 proteins in the non-smoking control group and 20 in the smoking group (Fig 1C). Additionally, 226 proteins were significantly upregulated and 136 were significantly downregulated (p value<0.05, FC ≥ 2) in HLM of heavy drinkers as compared to non-drinkers (Fig 1D). Overall, just 25 proteins showed significantly elevated levels in the HLM of moderate to heavy smokers (>1ppd) compared to non-smoking controls, while 62 proteins were significantly downregulated (p value<0.05, FC ≥ 1.5). Of note, numbers of significant changes in protein levels were reduced with lower levels of alcohol consumption. When protein expression in HLM from moderate drinkers was compared to non-drinkers, the number of proteins significantly upregulated was approximately 50% less and fewer than 15 were significantly downregulated, while less than 50 proteins had significantly altered levels in either the social or light drinker groups as compared to the non-drinking control group (Suppl Fig 1A). Input of significantly upregulated proteins from our dataset for STRING analysis revealed pathways involved in protein degradation, amino acid maintenance, and energy regulation being upregulated, while downregulated pathways included those involved in liver health and bile acid homeostasis, metabolism of complex carbohydrates, drug metabolism by P450s, cholesterol homeostasis, and oxidative phosphorylation (Suppl Fig 2).

### 3.2. Effects of alcohol intake on DMET proteome abundance and composition

To interrogate the impact of alcohol consumption on DMET protein expression, we assessed the absolute amounts of proteins based on the TPA approach from the HLM of each alcohol intake stratified group as compared to the non-drinking control group. A stark difference becomes apparent when evaluating the number of DMET proteins with significantly altered levels of expression in each alcohol intake category as compared to the non-drinking control (Fig 1F). Whereas significantly altered protein levels were observed for many DMET proteins in HLM from heavy drinkers as compared to non-drinking control, significant changes in the levels of DMET proteins were minimal in HLM of light, social, and moderate drinkers as compared to the non-drinking controls (Fig 1F, Suppl Fig 1B). For this reason, further analyses of the effects of alcohol on the liver proteome exclude light, social, and moderate alcohol intake groups. Overall, the majority of DMETs with significantly altered protein levels in HLM from heavy alcohol drinkers showed a decrease in expression as compared to non-drinkers, except for CYP2E1, UGT1A6, UGT1A9, FMO1, and ABCC3, which were elevated with heavy alcohol intake (p value<0.05, FC>1.25) (Fig 1G). The mean protein levels of CYPs, UGTs, and non-CYP, non-UGT drug metabolizing enzymes quantified in HLM from non-drinking control and the heavy alcohol intake group are reported with standard deviation and significance levels in Table 2.

**Table 2.**
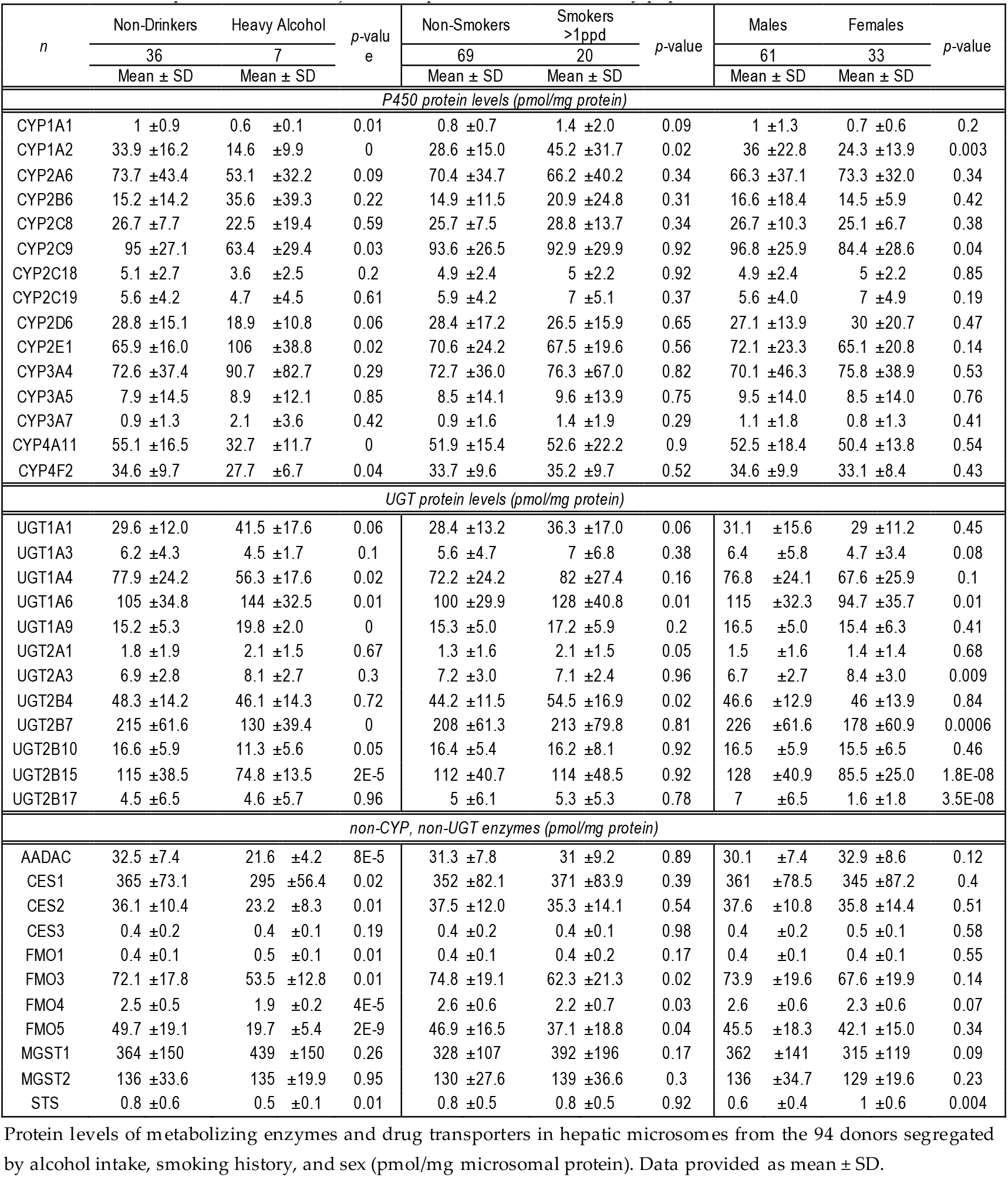
TPA based protein levels of major DMET proteins in HLM of study population.

Differential cytochrome P450 expression was seen in HLM of heavy drinkers as compared to non-drinkers (Fig 2A, Table 2). CYP1A1 (FC 0.57, p value <0.01), CYP1A2 (FC 0.43, p value<0.001), CYP2C9 (FC 0.67, p value<0.05), CYP4F2 (FC 0.8, p value<0.05), and CYP4A11 (FC 0.59, p value<0.001) protein levels were significantly reduced with heavy alcohol intake as compared to non-drinkers while CYP2E1 protein levels were significantly upregulated (FC 1.6, P<0.05). A decrease in the level of CYP2D6 approaching the level of significance was also observed (FC 0.65, P value=0.06) with heavy alcohol intake.

**Fig. 2.**
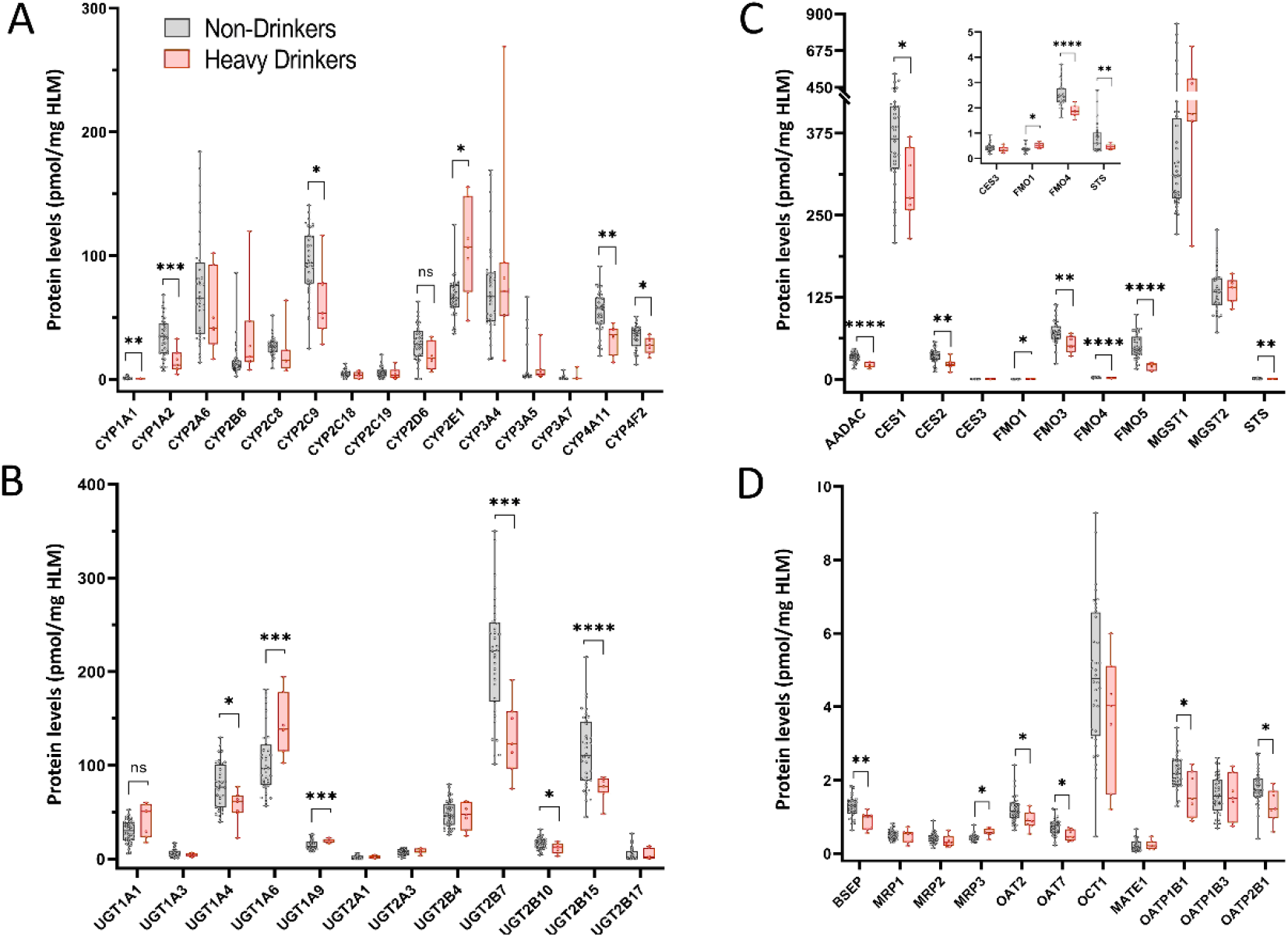
Differential DMET protein expression with heavy alcohol intake. A) Protein levels of major drug metabolizing CYPs in HLM from heavy drinkers compared to non-drinkers. B) UGT protein levels in HLM from heavy drinkers compared to non-drinkers. C) Protein levels of major non-CYP, non-UGT enzymes in HLM of individuals with heavy alcohol intake compared to non-drinkers. D) Transporter protein levels in HLM of heavy drinkers compared to non-drinkers. Samples were analyzed for significance using the student t-test with Welch’s correction for unequal variance. P Value *<0.05, **<0.01, ***<0.001, ****<0.0001. Absolute protein concentration for each DMET protein, expressed as pmol/mg of total HLM protein, was calculated via the TPA method.

As illustrated in Fig. 3A, these significant changes resulted in a considerable alteration of the composition of the P450 pool. Thus, the fraction of CYP2E1 increased from 11.4% in non-drinkers to 20% in heavy drinkers, while CYP1A1 and CYP1A2 decreased from 0.2 to 0.1 % and 5.9 to 2.8 %, respectively. CYP2C9 was significantly less abundant shifting down to 12% of overall CYP protein composition in heavy drinkers from 16.4% in non-drinkers, while CYP4A11 and CYP4F2 were reduced from 9.5 and 6% in non-drinkers to 6.2 and 5.2% in heavy drinkers. CYP4F11, CYP4F3, and CYP8B1 underwent a slight but statistically significant reduction with heavy alcohol intake, shifting from 1.8 to 1.7%, 2.2 to 1.8%, and 5.6 to 4.6%, respectively.

**Fig. 3.**
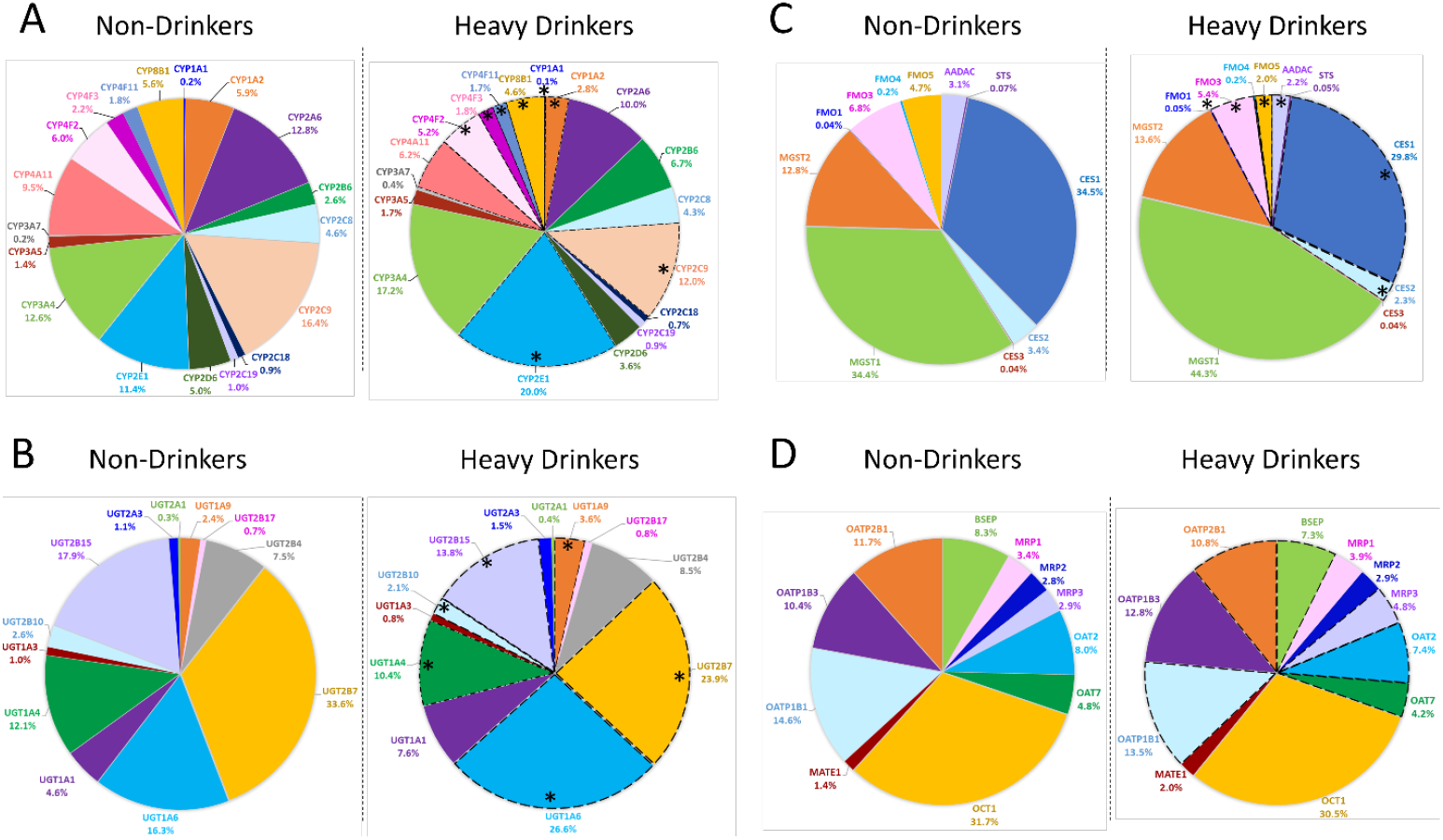
Shifts in overall composition of proteins related to drug metabolism and disposition with excessive alcohol intake. A) CYP protein composition in HLM from heavy alcohol drinkers compared to non-drinkers. B) UGT protein composition in HLM of heavy drinkers as compared to non-drinkers C) Non-CYP, non-UGT enzyme composition in HLM of heavy drinkers compared to HLM from non-drinkers. D) Composition of transporters measured in HLM of heavy alcohol drinkers as compared to that of non-drinkers. Absolute protein concentration for each protein was calculated using the TPA method and samples were analyzed for significance using the student t-test with Welch’s correction for unequal variance; * P value <0.05 indicates significant changes in protein levels were observed in the HLM of individuals with heavy alcohol intake as compared to that of the non-drinking control group.

Heavy alcohol consumption also significantly altered the levels of UGTs (Fig 2B, Table 2). Protein levels were significantly lowered for UGT1A4 (FC 0.7, p value<0.05), UGT2B7 (FC 0.6, p value<0.001), UGT2B10 (FC 0.68, p value<0.05), and UGT2B15 (FC 0.65, p value<0.0001), while the levels of UGT1A6 (FC 1.4, p value<0.01) and UGT1A9 (FC 1.3, p value<0.001) were significantly elevated. A moderate reduction in UGT1A1 approaching significance (FC 1.4, p value=0.06) was also observed.

The alcohol-induced changes in UGT abundances described above resulted in significant alteration of the composition of UGT pool (Fig 3B). In non-drinkers, UGT2B7 comprised almost 34% of all UGT enzymes expressed, while UGT2B15 made up 18%, UGT1A6 16%, and UGT1A4 12% of UGT abundance. In heavy drinkers the protein composition shifted significantly, such that the UGT1A6 fraction increased by 11% to 27% and the content of UGT2B7 decreased by 10% to 24%, while UGT2B15 and UGT1A4 were reduced to 13.8% and 10.4%, respectively. Additionally, UGT1A9 was increased from 2.4% in non-drinkers to 3.6% in heavy drinkers.

The expression pattern of non-CYP and non-UGT enzymes was also considerably altered by heavy alcohol intake (Fig 2C, Table 2). In particular, protein levels of AADAC (FC 0.66, p value<0.0001), CES1 (FC 0.81, p value<0.05) CES2 (FC 0.64, p value<0.01), FMO3 (FC 0.74, p value<0.01), FMO4 (FC .74, p value<0.0001), FMO5 (FC 0.4, p value<0.0001), and STS (FC 0.61, p value<0.01) were reduced, while, FMO1 (FC 1.36, p value<0.05) levels were elevated.

Significant changes were also observed in the levels of transporter proteins quantified in the HLM of heavy drinkers (Fig 2D). While MRP3 protein levels were significantly elevated (FC 1.26, p value<0.05), levels of BSEP (FC 0.7, p value<0.01), OAT2 (FC 0.73, p value<0.05), OAT7 (FC 0.68, p value<0.01), OATP1B1 (FC 0.68, p value<0.05), and OATP2B1 (FC 0.71, p value<0.05) were decreased.

The composition of non-CYP, non-UGT enzyme composition was altered with heavy alcohol intake (Fig 3C). We identified a significant decrease in the fractional content of AADAC, which dropped from 3.1% in non-drinkers to 2.2% in heavy alcohol consumers. CES1 and STS protein levels decreased from 34.5% to 29.8% and 0.07 to 0.05% (a 1.4-fold decrease), respectively. The levels of FMO3, FMO4, and FMO5 decreased from 6.8 to 5.4%, from 0.24 to 0.19%, and from 4.7 to 2%, respectively. In contrast, FMO1 protein levels increased with heavy alcohol consumption from 0.04% to 0.05% (a 1.25-fold increase).

The composition of transporters in HLM of heavy alcohol drinkers was also altered as compared to non-drinkers (Fig 3D). BSEP, OATP1B1, and OATP2B1 protein levels were significantly altered in overall composition, decreasing from 8.3 to 7.3%, 14.6 to 13.5% and 11.7 to 1.8%, respectively. Protein expression of OAT2 and OAT7 was significantly decreased and thereby lessened from 8 to 7.4% and 4.8 to 4.2%, respectively, in HLM from heavy drinkers as compared to non-drinkers. Meanwhile, MRP3 protein levels increased significantly, comprising almost 5% of transporters detected in HLM of heavy drinkers as compared to 2.9% in non-drinkers.

Additionally, further analysis was performed to account for smoking as a potential confounding factor by excluding specimens from individuals who smoked greater than 1 pack of cigarettes per day (Suppl Table 2). Though this test suffered from a lack of power, the trends in altered patterns of DMET protein expression levels remained the same, except for CYP2C8 which showed a significant decrease in protein levels in HLM from non-smoking heavy drinkers.

### 3.3. Establishing a provisional index of alcohol exposure based on the abundances of marker proteins and its use for in-depth analysis of the alcohol effects on HLM proteome

Although the analysis of differences between the four groups of donors described above provided valuable results on the effects of alcohol on the HLM proteome, this approach suffers from roughness due to voluntary and approximate reporting of the level of alcohol consumption by the liver donors. To improve the gradation of the liver samples and make our correlational analysis more robust, we sought to establish a scale of alcohol consumption based on the abundance of the protein markers of alcohol exposure in HLM.

For the analysis of correlations between protein abundances and the level of alcohol exposure of HLM donors, we selected a set of 75 proteins with known localization in the microsomal membrane or lumen (see Table 1 in Supplementary Materials for the list). Besides cytochromes P450 and their interaction partners (cytochrome b_5_, CPR, PGRMC1, and heme oxygenases), the set also included other microsomal drug-metabolizing enzymes (UGTs, FMOs, glutathione S-transferases, esterases CES2 and CES3). It also comprised the proteins involved in cellular response to ER stress (chaperones HSPA5, HSPA9, and HSPA90B1, protein disulfide isomerases, ER oxidoreductases ERO1A and ERO1B, and transitional ER ATPase (VCP) and some other relevant proteins (PGRMC2, STS, AADAC, ABCB6).

The MS intensities obtained in proteomic quantitation of each of these proteins were used to obtain the vectors of relative abundance (VRA) reflecting the relative differences in protein abundances between the HLM samples (see Materials and Methods). These vectors were then probed for their correlations with the vector of apparent alcohol consumption grade (ACG) of HLM donors. To build the ACG vector we used a provisional scale, where the donors indicated as “non-drinkers”, “former drinkers”, “light drinkers”, “social drinkers”, “low-to-moderate drinkers”, “moderate drinkers” and “heavy drinkers” were assigned the grades of 0, 0.5, 1, 1.5, 2, 3 and 4, respectively (see Materials and Methods).

Initially, we probed the correlation of ACG with the abundance vectors of every individual protein in the set. The best correlation (R^2^=0.23) was found with the abundance of the flavin-containing monooxygenase FMO5. This protein was followed with R^2^=0.215 by the heat shock protein HSPA5 (glucose-responsive protein 78, GRP78), a chaperone involved in the cellular response to the ER stress, and the transitional ER ATPase (VCP, R^2^=0.21), a protein responsible for exporting misfolded proteins from ER to cytoplasm. While the effects of alcohol on the abundances of VCP and FMO5 were not yet reported, the alcohol-induced increase in the expression of GRP78 is well established (Miles, 1995; Tunici et al., 1999; Rahman and Miles, 2001).

To better approximate the ACG vector, we then probed every possible combination of up to four proteins in the set. In this search, we approximated the ACG vector by a linear combination of several VRA using the multidimensional linear least-square regression algorithm (see Materials and Methods). This algorithm was sequentially applied to every possible combination of 2 - 4 proteins from our set of 75 selected proteins. The best two-protein combination comprises HSPA5 and PDIA3 (R^2^=0.37). The best three-protein combination is HSPA5, PDIA3, and CES2 (R^2^=0.44). The best four-protein combination includes the same proteins complemented with P4HB (R^2^=0.50). The multiplication coefficients for VRAs of HSPA5, P4HB, PDIA3, and CES2 in this combination are equal to 5.06, 2.35, -7.8, and 1.19.

The found combination of proteins contains three known markers of ER stress – endoplasmic reticulum chaperone HSPA5 and two protein disulfide isomerases – P4HB and PDIA3. Notably, despite the widely recognized role of these proteins in the cellular response to ER stress, no correlation of their abundance with alcohol exposure has been demonstrated so far. Interestingly, the two disulfide isomerases in this combination are taken with opposite signs – the contribution of P4HB is positive, whereas the one of PDIA3 is negative. Most likely, these opposite signs of contributions of two PDIAs are needed to distinguish alcohol exposure from other possible ER Stress inducers.

The correlation of this combination of four VRAs, which we termed the Provisional Index of Alcohol Exposure (PIAE), with the ACG is illustrated in Fig. 4. In this figure, panel A shows the correlation of PIAE with ACG of 94 individual HLM samples sorted in the order of increasing abundance of GRP78 (HSPA5), a known marker of alcohol exposure. The plot of PIAE versus ACG for the same HLM set is shown in Fig 4B. The correlation of PIAE with the reported level of alcohol consumption is well pronounced. Applying the Student’s T-test to probe the hypothesis of lack of correlation between these parameters results in a P-value as low as 3×10^-15^. Thus, PIAE can be used as a base for probing the correlation of relative abundances of microsomal proteins with the level of alcohol exposure of HLM donors. Results of this correlational analysis are shown in Table 3, which exemplifies the microsomal proteins exhibiting the T-test P-value below 0.05.

**Table 3.**
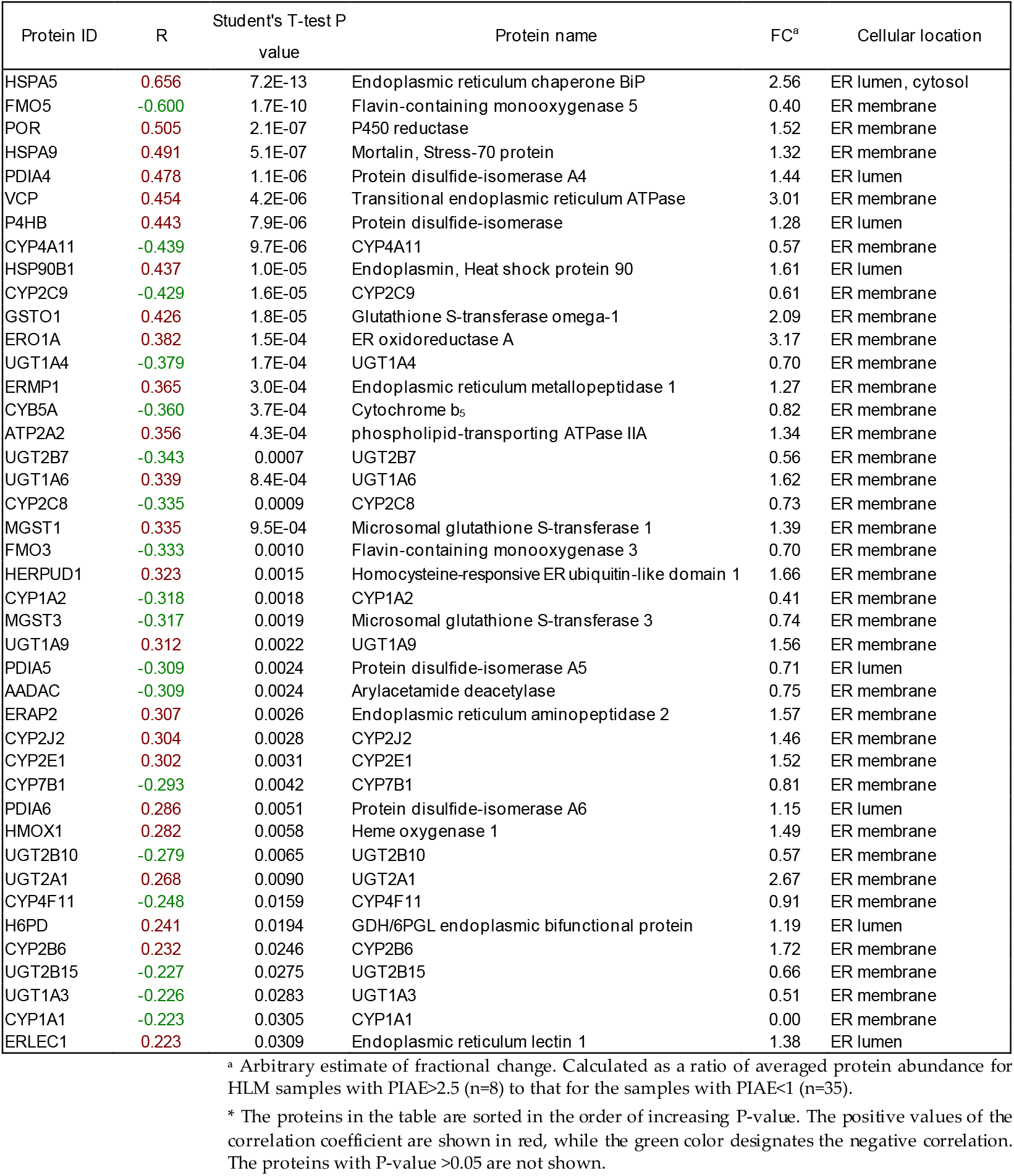
Microsomal proteins exhibiting a significant correlation between their relative abundance and the PIAE*.

**Fig. 4.**
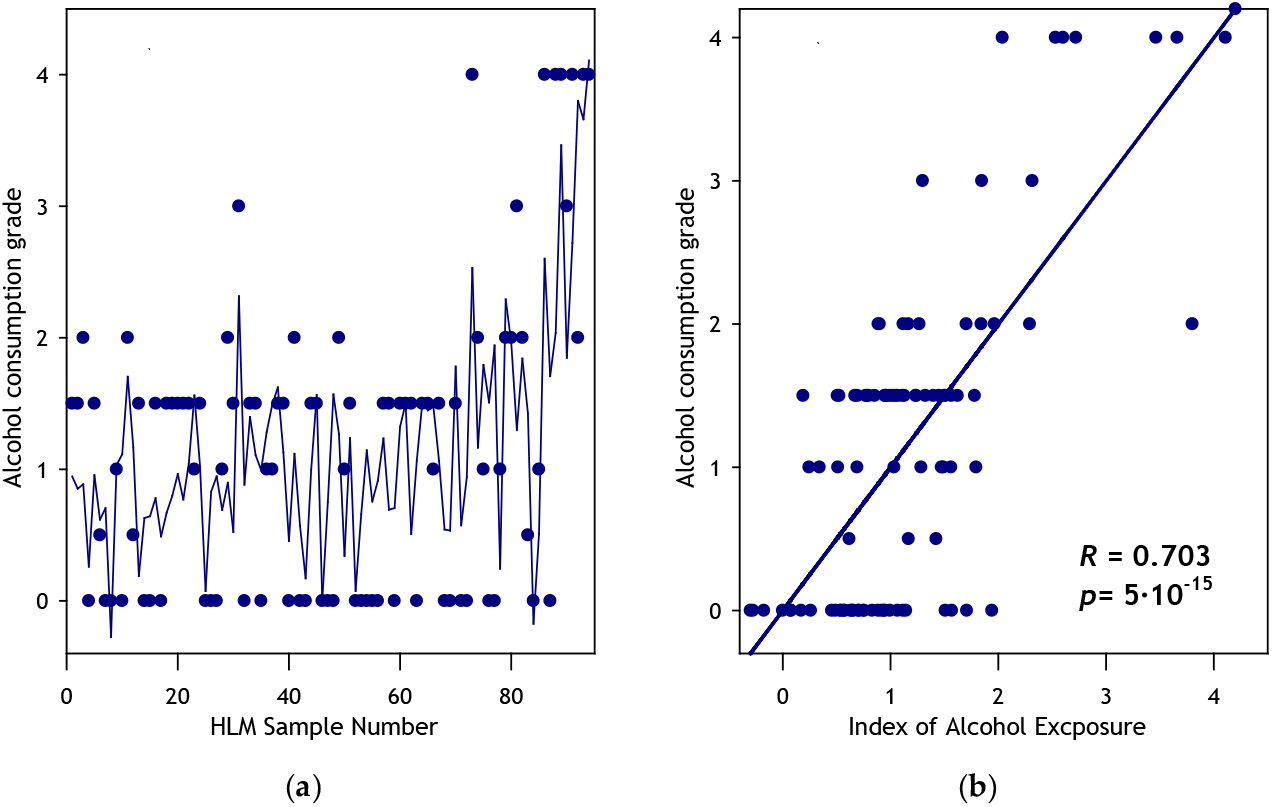
Approximation of the apparent alcohol exposure grade (ACG) with the provisional index of alcohol exposure (PIAE). The left panel shows the plots of the grade of alcohol exposure (circles) and the index of alcohol exposure (solid line) for all 94 HLM samples sorted by increasing the relative abundance of HSPA5 (GRP78). The right panel shows the same data as a plot of ACG versus PIAE.

As seen from this table, although many of the correlations picked up in our preliminary analysis were confirmed by analyzing the correlations with PIAE, there are also several notable differences between the results of the two approaches. In addition to UGT1A6 and UGT1A9 identified as upregulated by the preliminary analysis, we detected statistically significant upregulation of UGT2A1. The PIAE-based analysis also added UGT1A3 to the list of downregulated proteins while confirming the downregulation of UGT1A4, UGT2B7, and UGT2B15. Besides the upregulation of CYP2E1 detected by both methods, PIAE analysis also identified statistically significant upregulation of CYP2J2 and CYP2B6. Importantly, the PIAE-based analysis also identified a significant (p<10^-8^) upregulation of NADPH-cytochrome P450 reductase (POR ). While confirming the downregulation of CYP1A1, CYP1A2, CYP2C9, and CYP4A11, the PIAE analysis failed to confirm downregulation of CYP4F2 but complemented the list of downregulated proteins with CYP2C8, CYP4F11, and CYP7B1. It also indicates an alcohol-induced decrease in the abundance of cytochrome *b*_5_ (CYB5A, p<10^-4^) heme oxygenase 1 (HMOX1, p<10^-3^), and microsomal glutathione S-transferases MGST1 and MGST3. Table 4 summarizes the results of identifying the alcohol-induced changes in the abundances of UGTs, cytochromes P450, and their redox partners in HLM by PIAE analysis.

**Table 4.** Summary of the effects of alcohol exposure on the expression of drug-metabolized enzymes identified by PIAE analysis^*^.

| Protein class | Upregulated proteins | Downregulated proteins |
| --- | --- | --- |
| Cytochromes P450 | CYP2B6, <u>CYP2E1</u>,<br>CYP2J2 | <u>CYP1A1</u>, <u>CYP1A2</u>, CYP2C8, <u>CYP2C9</u>,<br><u>CYP4A11</u>, CYP4F11, CYP7B1 |
| Cytochrome P450 partners | POR, HMOX1 | CYB5A |
| UGTs | <u>UGT1A6</u>, <u>UGT1A9</u>,<br>UGT2A1 | UGT1A3, <u>UGT1A4</u>, <u>UGT2B7</u>, <u>UGT2B10</u>,<br><u>UGT2B15</u> |
| Other DMEs | GSTO1 | <u>AADAC</u>, <u>FMO3</u>, <u>FMO5</u>, MGST1, MGST3 |

\*The identifiers of the proteins revealed in our preliminary analysis are underlined.

### 3.4. Effects of Tobacco Smoke on DMET Abundance

Absolute protein concentration for each DMET protein, expressed as pmol/mg of total HLM protein, was calculated using the TPA method to assess the impact of moderate-to-heavy levels of smoking (>1 pack of cigarettes per day, or ppd) on DMET protein expression. In HLM from >1ppd tobacco users compared to non-smokers, CYP1A2 protein was upregulated significantly (FC 1.6, p value<0.05) and CYP1A1 protein levels were trending higher (FC 1.8, p value=0.09) with tobacco use (Fig 5A, Table 2). Expression of several UGTs was increased at the protein level in HLM from >1ppd tobacco users compared to non-smokers, specifically, UGT1A6 (FC 1.28, p value<0.05), UGT2A1 (FC 1.6, p value<0.05), and UGT2B4 (1.23, p value<0.05). Additionally, UGT1A1 protein was elevated (FC 1.28, p value=0.06) and approaching significance in tobacco users (Fig 5B, Table 2). Expression patterns of non-CYP, non-UGT enzyme levels in HLM of >1ppd smokers compared to non-smokers remained relatively stable with a few exceptions. In particular, significant decreases in the protein levels of FMO3 (FC 0.83, p value<0.05), FMO4 (FC 0.84, p value<0.05), and FMO5 (FC 0.79, p value<0.05) were observed in >1ppd smokers as compared to non-smokers (Fig 5C, Table 2). Transporter proteins quantified in HLM of >1ppd tobacco users compared to non-smokers show a significant decrease in protein levels of OAT7 (FC 0.83, p value<0.05) and OATP1B1 (FC 0.79, p value<0.05) with smoking (Fig 5D). When analysis was performed to account for potential confounding effects of alcohol by removing all heavy drinkers, similar trends in the levels of DMET proteins were seen (Supplementary Table 2).

**Fig. 5.**
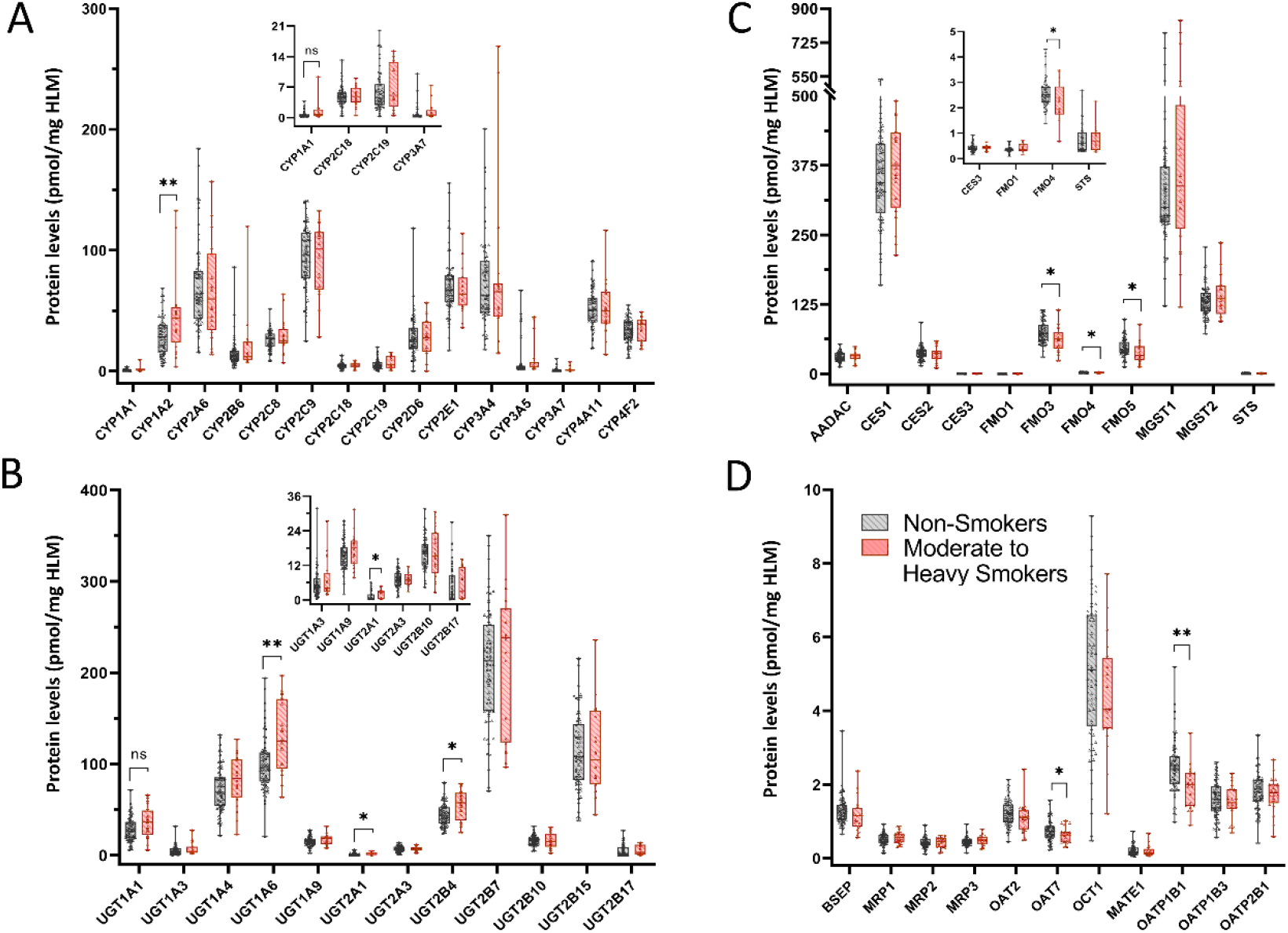
Differential DMET protein expression with moderate to heavy tobacco use. A) Protein levels of major drug metabolizing CYPs in HLM from >1 ppd smokers compared to non-smokers. B) UGT protein levels in HLM from >1 ppd smokers compared to non-smokers. C) Protein levels of major non-CYP, non-UGT enzymes in HLM of >1 ppd smokers compared to non-smokers. D) Transporter protein levels in HLM of >1 ppd smokers compared to non-smokers. Samples were analyzed for significance using the student t-test with Welch’s correction for unequal variance. P Value *<0.05, **<0.01. Absolute protein concentration for each DMET protein, expressed as pmol/mg of total HLM protein, was calculated via the TPA method.

### 3.5. Effects of Sex on DMET Abundance

Comparing the expression of CYPs in HLM from males to that of females, we observed significantly lower CYP1A2 (FC 0.7, p value<0.01) and CYP2C9 (FC 0.87, p value<0.05) protein levels in HLM from females (Fig 6A). Several UGTs had decreased protein levels in HLM from females, including UGT1A6 (FC 0.83, p value<0.05), UGT2B7 (FC 0.79, p value<0.001), UGT2B15 (FC 0.67, p value<0.0001), UGT2B17 (FC 0.2, p value<0.0001) as compared to males (Fig 6B, Table 2). Conversely, UGT2A3 protein levels were significantly elevated in females as compared to males. Protein levels of UGT1A3 were less in females when compared to males and approaching significance (FC 0.74), p value=0.079). Expression patterns of non-CYP, non-UGT enzyme levels in HLM of females remained unchanged when compared to males (Fig 6C, Table 2), except for STS, which was significantly elevated at the protein level in females (FC 1.55, p value<0.01). Similarly, transporter protein levels in HLM of females were comparable to that of males except for MRP1 (FC 0.8, p value=0.001), which was significantly reduced in HLM from females compared to males (Fig 6D, Table 2).

**Fig. 6.**
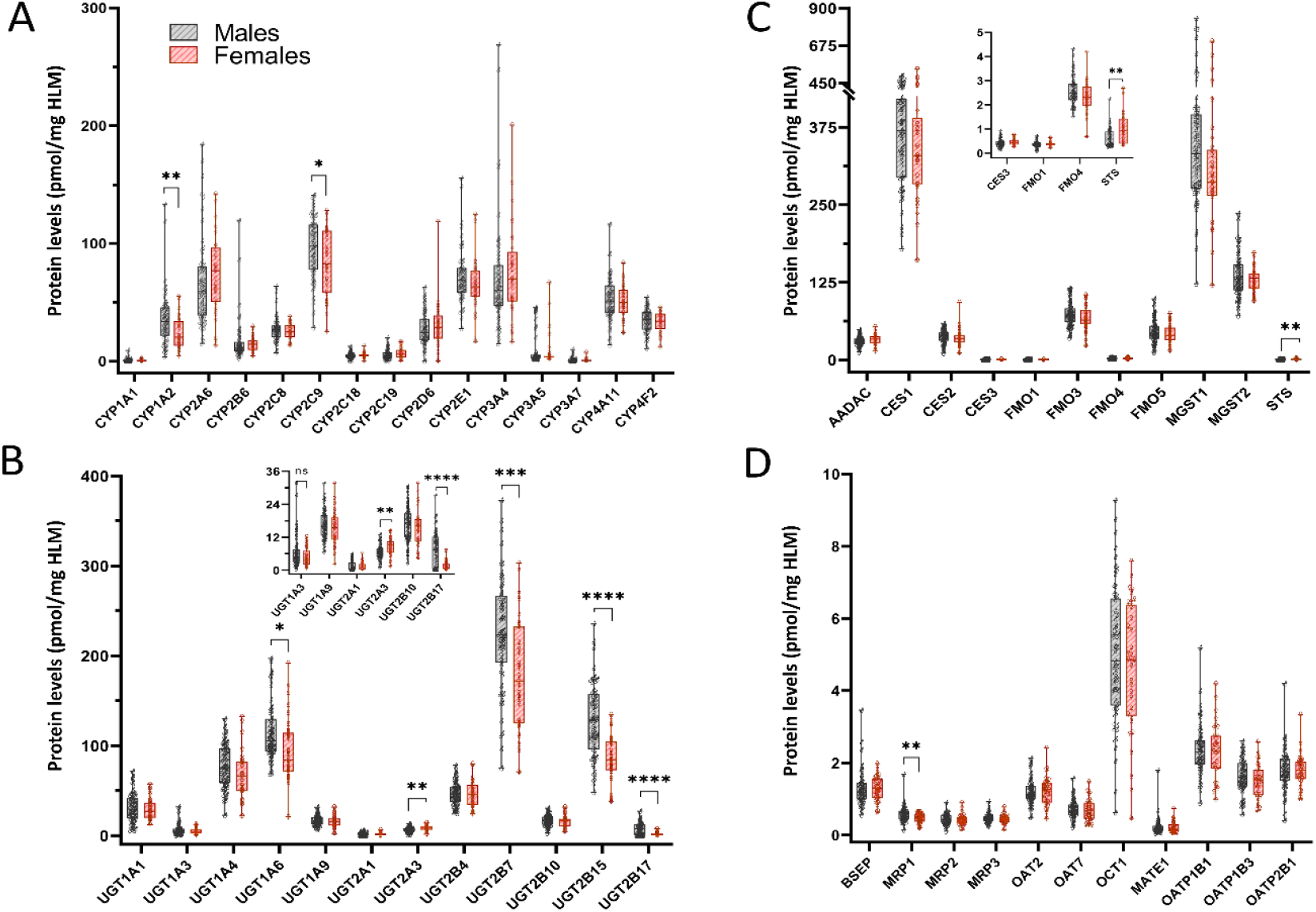
Differential DMET protein expression with sex. A) Protein levels of major drug metabolizing CYPs in HLM from female donors compared to males. B) UGT protein levels in HLM from female compared to male donors. C) Protein levels of major non-CYP, non-UGT enzymes in HLM female donors compared to males. D) Transporter protein levels in HLM females compared to males. Samples were analyzed for significance using the student t-test with Welch’s correction for unequal variance. P Value *<0.05, **<0.01, ***<0.001, ****<0.0001. Absolute protein concentration for each DMET protein, expressed as pmol/mg of total HLM protein, was calculated via the TPA method.

## 4. Discussion

Alcohol and tobacco use are prevalent and result in a wide range of health effects. While much is understood about tobacco metabolism, tobacco smoke is comprised of over 7000 chemicals, complicating our understanding of the potential for drug interactions. Additionally, knowledge of the effects of alcohol on the abundance of DMET proteins remains limited. While several studies have been carried out to examine the mechanisms of the effects of alcohol exposure on drug metabolism, investigations have been based on piecemeal examination of a few enzymes at a time in a variety of *in vitro* and *in vivo* models, and evidence is often conflicting. However, the ability to accurately assess potential drug-alcohol interactions in a systematic way is critical to establishing a safe and effective drug profile in the overall population (Neuhoff et al., 2021; Ahire et al., 2022; Basit et al., 2022).

In this study we used the total proteomics approach for comparing the abundance and composition of DMET proteins in a set of 94 HLM preparations from donors with documented history of alcohol consumption and tobacco to explore the effect of these factors on the composition of the human drug metabolizing system. We detected over 4000 proteins in our HLM samples overall and analyzed DMET protein levels in relation to alcohol consumption grades, smoking history data, and sex using non-parametric tests (p-value ≤ 0.05; cutoff of 1.25-fold change, FC). The examination of the alcohol-induced changes was further enforced by correlational analysis, where we used arbitrary alcohol consumption grade (ACG) scaling from 0 to 4 to establish a set of protein markers. We elaborated a provisional index of alcohol exposure (PIAE) based on a combination of relative abundances of four proteins best correlating with ACG. The set of found marker proteins is comprised of HSPA5, PDIA3, P4HB, and CES2. The PIAE index was then used to find its correlations with the abundances of DMET proteins. This innovative approach allowed us to corroborate and further extend the conclusions driven from the initial non-parametric analysis.

Our results demonstrate considerable alcohol-induced changes in the composition of the cytochrome P450 pool in HLM. Consistent with the literature (Cederbaum, 1998; Dupont et al., 1998; Lieber, 1999; Meskar et al., 2001; Cederbaum, 2006), we observed significant upregulation of CYP2E1 with alcohol consumption, which increased by 1.6-fold on average in heavy drinkers as compared to non-drinkers. Besides this well-known effect of alcohol, our correlational analysis reveals significant increases in the abundances of CYP2B6, CYP2J2, and NADPH-cytochrome P450 reductase (POR), which have never been reported in literature. The increase in the level of POR may, at least in some part, be responsible for the general increase in the rate of drug metabolism by alcohol consumption stated by several authors (Kater et al., 1969a; Kater et al., 1969b; Tanaka, 2003; Chan and Anderson, 2014). Importantly, we did not detect any alcohol-induced increase in the abundance of CYP3A4 or CYP2A6, which has been suggested in literature based on animal models and *in vitro* data (Feierman et al., 2003; Lu et al., 2011; Lu et al., 2012).

In addition to the above findings, our novel approach of combined proteomics analysis and use of the PIAE index to perform correlation analysis also uncovered a significant lowering of the levels of CYP1A1, CYP1A2, CYP2C8, CYP2C9, CYP4A11, CYP4F11, and CYP7B1 proteins by chronic alcohol exposure. Thus, the average levels of CYP1A2, CYP2C9, and CYP4A11 were decreased by 2 .3, 1.5, and 1.7-fold, respectively, with heavy alcohol consumption. Downregulation of CYP1A2, CYP2C8, and CYP2C9 by alcohol exposure may have a profound impact on pharmacokinetics of a variety of drugs metabolized by these enzymes. At the same time, an alcohol-induced decrease in the abundance of CYP4A11, the enzyme involved in the synthesis of eicosanoids, may affect the 20-HETE signaling pathways and be involved in the mechanisms of alcohol-induced hypertension. The observed decrease in the level of cytochrome *b*_5_ with alcohol exposure may also have important physiological consequences. Interactions of cytochrome *b*_5_ with cytochromes P450 are known to increase the coupling of the cytochrome P450 ensemble and reduce the P450-dependent production of harmful reactive oxygen species (Pompon, 1987; Perret and Pompon, 1998; Schenkman and Jansson, 1999). Thus, the alcohol-induced decrease in the level of cytochrome *b*_5_ in the ER membrane may be involved in the mechanisms of ethanol hepatotoxicity.

Considerable alcohol-induced changes in drug metabolism may also be caused by the changes in the abundance and composition of the ensemble of UGTs. We observed significant upregulation of UGTs 1A6, 1A9, and 2A1; and significant downregulation of UGTs 1A3, 1A4, 2B7, 2B10, and 2B15 with alcohol exposure. Protein levels of UGT1A6 and UGT1A9, which have been suggested to play a role in non-oxidative metabolism of ethanol, (Hugbart et al., 2020) were also significantly elevated (1.4 and 1.3-fold increase) in HLM of heavy drinkers versus non-drinkers, which has not been previously reported. When smokers were removed from the analysis, the effects of alcohol remained similar (Supplementary Table 2).

In HLM from moderate-to-heavy smokers (>1 pack a day; n=13) significant upregulation of CYP1A2 (FC 1.6), UGT1A6 (FC 1.28), UGT2A1 (FC 1.6), and UGT2B4 (1.23) and significant downregulation of FMO3 (FC 0.8), FMO4 (FC 0.8), and FMO5 (FC 0.8) was observed as compared to non-smokers (n=60), independent of alcohol consumption. While CYP1A2 is known to be elevated with smoking due to its role in detoxification of polyaromatic hydrocarbons, and there are reports in the literature of increased activity of UGT1A6 with cigarette smoke, population-based studies into potential induction of these enzymes in the liver due to smoking is lacking.

Male donors (n=52) showed significantly higher abundance of CYP1A2 (FC 1.4), UGT2B17 (FC 4.5), UGT2B15 (FC 1.5), and a significantly lower abundance of UGT2A3 (FC 0.75) as compared to females (n=30). While an association of UGT2B17 abundance with sex is well-known, the effect of sex on other DMET proteins of clinical importance is a novel finding.

Limitations due to the nature of the study only allow for observational analysis and do not allow for controlled environment. Therefore, it is possible that there were some confounding factors such as second-hand smoking which could have led to an elevation of CYP1A1 or CYP1A2 in some individual donors from the control group. However, there are also benefits to a population-based study, which include the realistic nature of the study design and ability to determine significant changes in spite of inter-individual variations.

Previously, we have shown evidence of an impact on CYP3A4, CYP1A2, and CYP2C19 activity due to induction of CYP2E1 by alcohol (Davydova et al., 2019; Dangi et al., 2021), indicating the importance of consideration of the mutual functional effects of multiple enzymes constituting thanalyzing the correlatione P450 ensemble. The effects of alcohol observed here on the induction or suppression of protein expression of different CYPs, therefore, could have additional impacts on the functioning of the drug-metabolizing ensemble as a whole. The use of the Provisional Index of Alcohol Exposure elaborated in this study for analyzing the correlations between the degree of alcohol exposure and the function of drug-metabolizing enzymes in HLM offers a prospective way for further exploration of the effects of alcohol on the drug-metabolizing system.

In summary, this represents the first comprehensive study to our knowledge, to report the effects of alcohol intake, smoking, and sex differences on the protein levels of important DMET proteins in a large set of human liver samples. Our data showed alcohol-associated differences in the abundance and composition of important DMET proteins involved in the metabolism and disposition of a variety of pharmaceuticals, as well as the need for smoking tobacco and sex to be considered as co-variates.

## Supplementary Materials

The following supporting information can be downloaded: Figure S1: Global and DMET proteome analysis of HLM from light, social, and moderate alcohol drinkers., Figure S2: STRING analysis of significantly upregulated and downregulated proteins from global proteomics, Table S1: List of proteins localized to the endoplasmic reticulum used for normalizing DMET proteins, Table S2: TPA based protein levels of major DMET proteins present in HLM of heavy drinkers with >1ppd smokers removed and smokers with heavy drinkers removed.

## Author Contributions

Conceptualization, B.P. and D.R.D.; methodology, B.P., D.R.D., and P.L.; software, D.D.; formal analysis, K.A.G., D.R.D., and B.P.; investigation, D.K.S., K.A.G., N.Y.D., and K.P.; resources, D.R.D., B.P., and P.L.; writing— K.A.G. and D.R.D.; writing – review and editing D.R.D. and B.P. All authors have read and agreed to the published version of the manuscript.

## Data Availability Statement

The data are contained within the article. The raw data sets used to generate the reported results are available from the authors upon a reasonable request.

