## Supplemental Figures 1 and 2 for "Effects of alcohol consumption and tobacco smoking on the composition of the ensemble of drug metabolizing enzymes and transporters in human liver"

### Slide 1
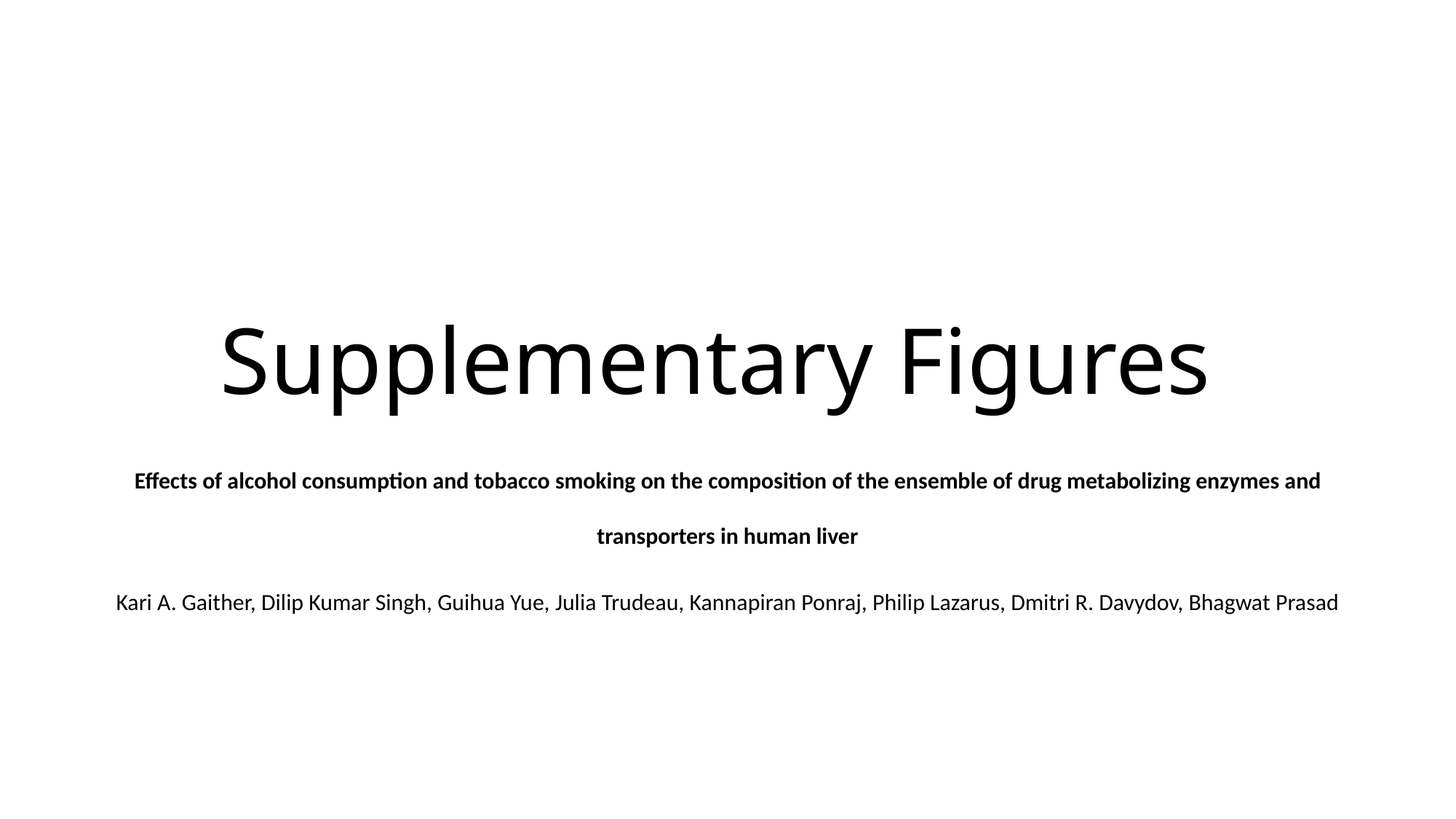

#### Supplementary Figures
Effects of alcohol consumption and tobacco smoking on the composition of the ensemble of drug metabolizing enzymes and transporters in human liver
Kari A. Gaither, Dilip Kumar Singh, Guihua Yue, Julia Trudeau, Kannapiran Ponraj, Philip Lazarus, Dmitri R. Davydov, Bhagwat Prasad

### Slide 2
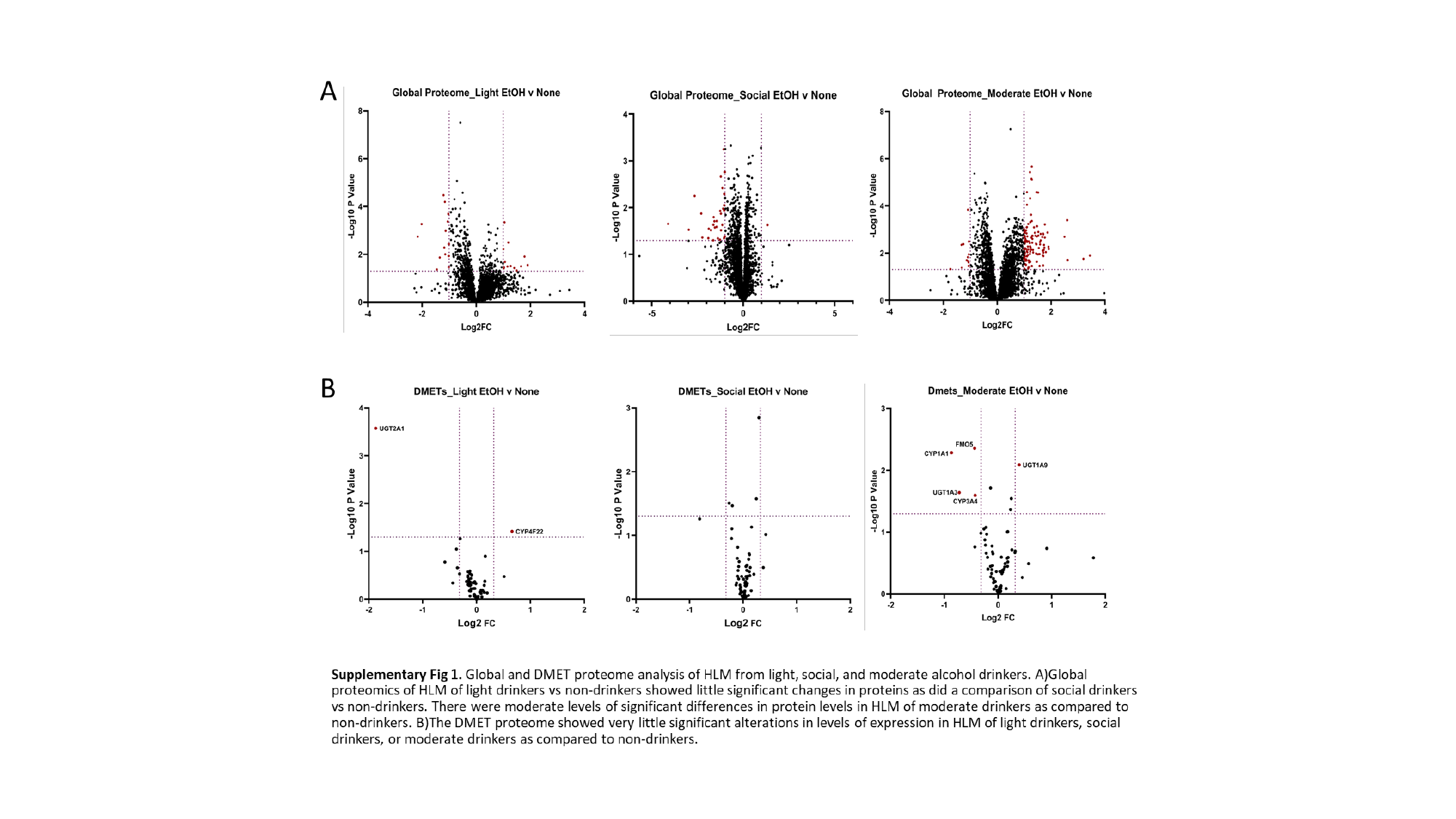

### Slide 3
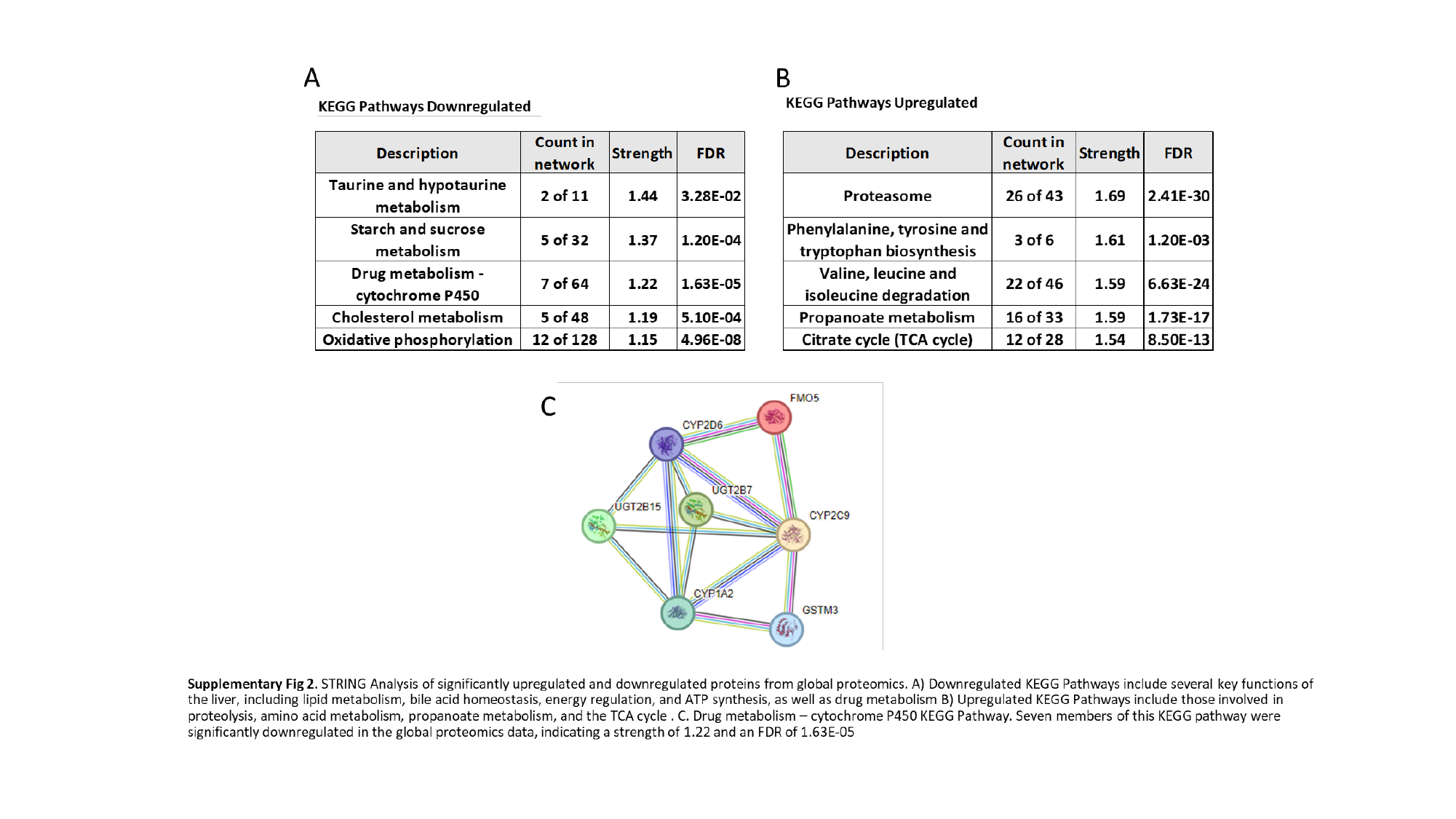
